# Epigenomic patterns reflect irrigation and grafting in the grapevine clone ‘Chambourcin’

**DOI:** 10.1101/2020.09.09.290072

**Authors:** Brigette R. Williams, Christine E. Edwards, Misha T. Kwasniewski, Allison J. Miller

**Affiliations:** Saint Louis University, Department of Biology, 3507 Laclede Ave, St. Louis, MO 63103, USA; Missouri Botanical Garden, Center for Conservation and Sustainable Development, 4651 Shaw Blvd., St. Louis, MO 63110, USA; The Pennsylvania State University, College of Agricultural Sciences Department of Food Science, 326 Rodney A. Erickson Food Science Building, University Park, PA 16802, USA; Donald Danforth Plant Science Center, 975 North Warson Road, St. Louis, MO 63132, USA

## Abstract

Although DNA methylation has largely been shown to be stable in plants, mounting evidence indicates methylation patterns may reflect environmental sensitivity. Perennial plants experience seasonal and inter-annual environmental variation, and clonal replicates of some long-lived plants, including many perennial crops, survive in a broad range of environments. This makes perennial crops a compelling study system to investigate links between the plant epigenome and environmental variation. In this study, we used whole genome bisulfite sequencing and small RNA sequencing to characterize the epigenome in 12 clonal replicates of the winegrape cultivar ‘Chambourcin.’ We asked whether DNA methylation varied in response to a full factorial combination of irrigation and grafting treatments. We found signatures of both irrigation and grafting in the ‘Chambourcin’ epigenome, as well as compelling evidence for a unique interaction effect whereby grafting appeared to override or mitigate epigenomic changes associated with irrigation in ungrafted vines. These findings indicate that the epigenome responds to environmental and agronomic manipulations, suggesting the epigenome might be a mechanism underlying how long-lived, clonal plants respond at the molecular level to their environment. Further research is needed to assess the potential relevance of variation in DNA methylation to plant form and function, and to address the implications of environmentally-inducible patterns of DNA methylation on the adaptive capacity of long-lived woody perennials in nature and under cultivation.

## Introduction

Sessile organisms must be capable of surviving a wide range of environmental conditions. One way this is accomplished is through phenotypic plasticity; the ability of a genetic individual to produce a range of phenotypes under different environmental conditions. Recent work suggests a link between phenotypic plasticity and the epigenome, which enables an individual to alter gene expression through non-genetic mechanisms (Kooke et al., 2015; Zhang et al., 2018b; Perrone and Martinelli, 2020). Moreover, accumulating evidence supports the epigenome as a type of environmental memory in plants, whereby environmental events are recorded through epigenomic modifications that the plant relies on for heightened sensitivity and faster response to such events in the future (known as priming; Lämke and Bäurle, 2017). However, relatively little is known about the capacity of a single individual to remodel patterns of epigenomic variation, particularly DNA methylation, in complex plant-environment interactions.

The plant epigenome is a dynamic regulatory system of the genome involving covalent modifications to DNA, DNA-associating proteins (histones and histone variants) and some RNAs, and the activity of non-coding RNAs that can affect gene expression without changing the DNA sequence (Du et al., 2015). The epigenome manages genome stability and accessibility by creating and remodeling chromatin structure (Vergara and Gutierrez, 2017). DNA methylation patterns within an individual(i.e., the methylome), the most well-studied component of the epigenome, involves the establishment, maintenance, and removal of a methyl group to the 5’ position on cytosines (Law and Jacobsen, 2011). DNA methylation in plants occurs in three sequence contexts (CpG, CHG, and CHH, where H represents A, T, or C), and much of it appears to be controlled genetically (Dubin et al., 2015; Kawakatsu et al., 2016), with a large proportion being stable regardless of environment (Becker et al., 2011; Schmitz et al., 2011; Seymour et al., 2014).

Patterns of DNA methylation can also vary independently from genetic variation (Klironomos et al., 2013; Cortijo et al., 2014; Kooke et al., 2015; van der Graaf et al., 2015; Wilschut et al., 2016) and can change in response to environmental variation. For example, genome-wide remodeling of DNA methylation has been documented in response to both biotic factors (infectious pathogens, herbivory, and plant-plant competition) and abiotic factors (light, temperature, nutrient availability, and precipitation) (Verhoeven et al., 2010a; Dowen et al., 2012; Dubin et al., 2015; Secco et al., 2015; Kawakatsu et al., 2016; Alonso et al., 2019; Thiebaut et al., 2019). Furthermore, the methylomes of some plants show signatures of local adaptation and geographic provenance (Platt et al., 2015; Kawakatsu et al., 2016; Wilschut et al., 2016; Busconi et al., 2018). Such patterns of environmentally induced DNA methylation have been proposed as a form of environmental memory because they can be stably transmitted from parent to offspring (Boyko et al., 2010; Bilichak et al., 2012). Epigenomic modifications have also been transmitted clonally among ramets of a genet (parent plant), and are thought to have an important role in facilitating the persistence of clonal plants and their ability to adapt to fluctuating environments (Rendina González et al., 2016).

One group in which the ability to respond to environmental variation is particularly important is long-lived (perennial), woody plants. Perennial, woody plants must withstand environmental variation occurring over days, weeks, years, decades, centuries (*Quercus rubra* L.), and even millennia (*Taxodium distichum*; Stahle et al., 2019). Phenotypic plasticity is an important mechanism enabling long-lived plants to respond to changes in environmental conditions. Epigenomic responses to environmental variation is both a form of phenotypic plasticity and a mechanism that may underlie phenotypic plasticity in other traits through its regulatory effects on the genome (Johnson and Tricker, 2010; Kooke et al., 2015). Despite the potential importance of epigenomic plasticity in the ability of long-lived, woody plants to respond to their environment, the specific environmental drivers associated with epigenomic plasticity, and the extent to which such plants demonstrate epigenomic plasticity are largely unknown. Most previous studies of epigenomic plasticity have been conducted in annual species, including the model system *Arabidopsis* (Johannes et al., 2009; Gao et al., 2010; Verhoeven et al., 2010; Paun et al., 2010; Dowen et al., 2012; Weigel and Colot, 2012; Stroud et al., 2013; Zhong et al., 2013; Herman and Sultan, 2016; Wilschut et al., 2016; Alvarez et al., 2018). Studies of epigenomic plasticity in non-model woody perennials have typically employed low-resolution methods like MSAP (methylation sensitive amplified polymorphism) to quantify and describe patterns of epigenetic variation in experimental conditions and natural settings (Lira-Medeiros et al., 2010; Herrera and Bazaga, 2011; Saéz-Laguna et al., 2014; Xie et al., 2017). These studies suggest some degree of environmental sensitivity in the plant epigenome. However, it is difficult to identify clear associations between changes in methylation and environmental differences in such studies because the MSAP method surveys anonymous loci, for which detection partially depends on conserved genetic sequences at restriction sites, rather than on the strict presence-absence of methylation. Therefore, more comprehensive assessments of whole genome methylation patterns are required to detect subtle changes associated with environmental influences.

An approach to quantify epigenetic variation that provides greater resolution of patterns of DNA methylation is bisulfite sequencing (BS-seq), where genomic DNA is subjected to high-throughput DNA sequencing after being treated with bisulfite, which converts unmethylated cytosines to uracil, so all remaining cytosines are assumed to be methylated. For example, Heer et al. (2018) used BS-seq to investigate the effects of the environment on methylation in Norway spruce (*Pinus abies*). In this study, four genotypes replicated at two distinct sites showed a strong signature of clonal genotype in patterns of DNA methylation but little effects of site (Heer et al 2018). Another study found bud break in poplar was controlled by an environmentally induced epigenomic mechanism, whereby low temperatures activated a demethylase that reduced methylation that then reactivated genes controlling the shift from winter dormancy to vegetative growth (Conde et al., 2018). The extent to which the environment affects the epigenome of plants, as well as which environmental conditions induce epigenomic plasticity, have important implications for understanding how long-lived plants respond to and persist through changes in their environment.

Clonal plants offer an ideal system for understanding the extent to which the epigenome responds to environmental variation while controlling for genetic variation. Important both ecologically and agriculturally, it is estimated that clones make up 40% of all plants (Tiffney et al., 1985) and more than 75% of woody crops are clonally propagated (Miller and Gross 2011), with some popular clonal varieties cultivated nearly worldwide. Epigenomic plasticity might be one mechanism that allows the same clonal genotype to thrive in a wide range of climates (Douhovnikoff and Dodd, 2014; Dodd and Douhovnikoff, 2016). Clonal plants provide a powerful experimental system in which clonal replicates of the same genotype can be subjected to environmental manipulations applied consistently over multiple years, offering an unusual opportunity to characterize epigenomic plasticity in response to the environment.

Domesticated at least 6,000 years ago, cultivated grapevines (*Vitis* spp.) are among the oldest clonally propagated perennial crops (Pinhas and Spiegl-Roy, 1975; This et al., 2006, Myles et al., 2011). Clonally propagated grapevine cultivars are grown across very different climates, where they have been shown to exhibit phenotypic variation (Dal Santo et al., 2013; Chitwood et al., 2016; Young et al., 2016). Many grapevine breeders employ grafting, a surgical enhancement that joins the aboveground portion of one plant (scion) to the root system of another individual (rootstock). Previous research has shown that both the environment and the rootstock have important effects on the phenotype of the grapevine scion (Dal Santo et al., 2013; Anesi et al., 2015; Chitwood et al., 2016b; Young et al., 2016; Migicovsky et al., 2019), and both have consequences for a wine’s *terroir* (the concept that a wine’s flavor is of a place, derived from a specific set of environmental, soil and microbial conditions that produce stark, but consistent flavor differences between clonally propagated cultivars grown in different regions) (Sabon et al., 2002). Furthermore, prior work also indicated that different clonal lineages of the wine grape cultivar Pinot noir can be distinguished by MSAP profiles (Ocaña et al., 2013). Additionally, Xie et al. (2017) detected a signature of geographic provenance in DNA methylation in 198 Shiraz vines sampled from 22 vineyards across Australia, suggesting that the epigenome might encode a vine’s *terroir*. However, the extent to which the environment can influence variation in DNA methylation in grapevine clones is currently unknown.

Using whole genome BS-seq in 12 replicates of a single clonal grapevine cultivar ‘Chambourcin,’ we investigated how patterns of DNA methylation vary in an individual genotype under different irrigation treatments, when grafted, and the interaction of these conditions. In addition, in a subset of eight of the 12 replicates, we sequenced and measured the abundance of small RNAs, which are responsible for *de novo* establishment and maintenance of DNA methylation through the RNA-directed DNA methylation pathway (RdDM; Matzke and Mosher, 2014; Movahedi et al., 2015; Springer et al., 2015; Lewsey et al., 2016; Bouyer et al., 2017; Tamiru et al., 2018). We addressed the following questions: (1) How does irrigation affect the methylome? (2) How does grafting affect the methylome? And (3) How does the interaction of irrigation and grafting affect the methylome? To our knowledge, this study is among the first to document epigenomic plasticity in a clonal woody individual that was subjected to multi-year experimental treatments in the field, providing information on one mechanism that might underlie plasticity in long-lived clonal plants. Furthermore, by investigating the long-term effects of environment on the epigenome of clonally propagated grapevine scions, this work has potential implications for explaining an element of the viticultural concept of *terroir*.

## Results

### Whole Genome Bisulfite Sequencing

#### Genome mapping of BS-seq reads

We used whole genome BS-seq to investigate cytosine methylation among 12 clonal replicates of the winegrape cultivar ‘Chambourcin’ in an experimental vineyard at the University of Missouri Southwest Research Center (Mount Vernon, MO, USA). The 12 ‘Chambourcin’ replicates were divided evenly among four experimental treatments (Fig. 1A): Ungrafted, Unirrigated (UG-UI); Ungrafted, Irrigated (UG-IR); Grafted, Unirrigated (GR-UI); and Grafted, Irrigated (GR-IR). Grafted vines consisted of ‘Chambourcin’ scions grafted to the rootstock cultivar ‘3309C.’ Ungrafted vines are ‘Chambourcin’ growing ungrafted on their own roots. The irrigation treatments were initiated in 2015 after vines were established; UI vines received no irrigation from 2015 onwards and IR vines received full replacement of evapotranspiration. Leaf tissue was collected on September 25, 2017, immediately preceding harvest, placed in cryo-vials in liquid nitrogen in the field, and stored at −80C in the lab. Methylome profiles were generated for each of the 12 individuals. Whole-genome BS-seq reads were mapped to the *Vitis vinifera* reference genome (PN40024 12Xv3). Total coverage depth of the genome was approximately 80x, with coverage depth per replicate group estimated to be 20x (Table 1). Forty-three-point-four −50.5% of bisulfite converted reads mapped to unique locations in the genome (see Table S1); 12.1 – 14.0% of reads multi-mapped. Mapping bisulfite-converted reads against the organism’s chloroplast genome is an efficient and accurate method for estimating bisulfite conversion rate because chloroplast genomes are typically unmethylated (Feng et al., 2010). We mapped BS-seq reads against the *Vitis* chloroplast genome (NCBI GenBank RefSeq: NC_007957) and estimated the mean bisulfite conversion rate at 97.7% (see Table S1).

**Figure 1.**
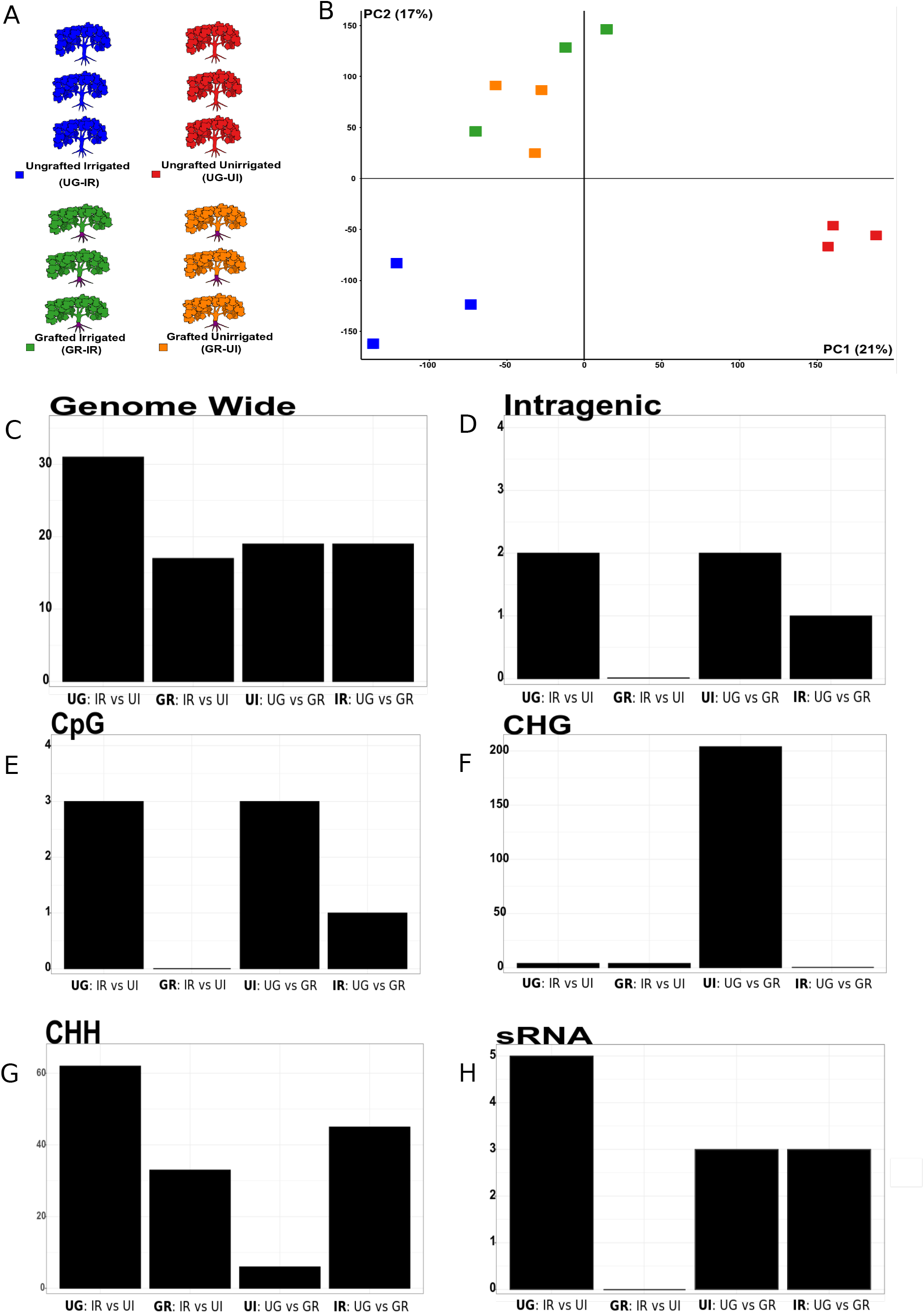
Patterns of variation in DNA methylation and sRNA among experimental groups. **(A)** Experimental design of replicate vines. Ungrafted-Irrigated (UG-IR; blue). Ungrafted-Unirrigated (UG-UI; red). Grafted-Irrigated (GR-IR; green). Grafted-Unirrigated (GR-UI; orange). Vines are planted and irrigated in a 2×2 factorial design (factors: irrigation, grafting). Sampled vines are located in the center of the vineyard and three clonal replicate vines per treatment block were sampled for a total of 12 clones included in this study. See also Table S1. **(B)** Clonal replicate grapevines show distinct patterns of genome-wide DNA methylation by environment and rootstock. PCA of patterns of genome-wide cytosine methylation with ≥20% difference in methylation among all individual vines. PC1 (21%) suggests irrigation treatment is the primary factor underlying variation in differential DNA methylation. PC2 indicates root identity (or the condition of being grafted) is a secondary factor of differences in DNA methylation. This analysis demonstrates that ‘Chambourcin’ grapevines can be distinguished by signatures of irrigation and root identity in patterns of DNA methylation, and the condition of being grafted diminishes a signal of irrigation in the ‘Chambourcin’ methylome. See also Table S2. **(C)** genome-wide (in all sequence contexts) DMR counts for each experimental group comparison: 31 - Ungrafted, Irrigated vs Unirrigated (left); 17 - Grafted, Irrigated vs Unirrigated (center-left); 19 - Unirrigated, Ungrafted vs Grafted (center-right); 19 - Irrigated, Ungrafted vs Grafted (right). See also Table S3. **(D)** Intragenic (+/−2.5kb; all sequence contexts) DMR counts by experimental group comparison: 2 - Ungrafted, Irrigated vs Unirrigated (left); 0 - Grafted, Irrigated vs Unirrigated (center-left); 2 - Unirrigated, Ungrafted vs Grafted (center-right); 1 - Irrigated, Ungrafted vs Grafted (right). See also Table S3. **(E)** CpG sequence context only DMR counts by experimental group comparison: 3 - Ungrafted, Irrigated vs Unirrigated (left); 0 - Grafted, Irrigated vs Unirrigated (center-left); 3 - Unirrigated, Ungrafted vs Grafted (center-right); 1 - Irrigated, Ungrafted vs Grafted (right). See also Table S3. **(F)** CHG sequence context only DMR counts by experimental group comparison: 3 - Ungrafted, Irrigated vs Unirrigated (left); 4 - Grafted, Irrigated vs Unirrigated (center-left); 204 - Unirrigated, Ungrafted vs Grafted (center-right); 0 - Irrigated, Ungrafted vs Grafted (right). See also Table S3. **(G)** CHH sequence context only DMR counts by experimental group comparison: 61 - Ungrafted, Irrigated vs Unirrigated (left); 33 - Grafted, Irrigated vs Unirrigated (center-left); 6 - Unirrigated, Ungrafted vs Grafted (center-right); 45 - Irrigated, Ungrafted vs Grafted (right). See also Table S3. **(H)** DE sRNA counts by experimental group comparison: 5 - Ungrafted, Irrigated vs Unirrigated (left); 0 Grafted, Irrigated vs Unirrigated (center-left); 3 - Unirrigated, Ungrafted vs Grafted (center-right); 3 - Irrigated, Ungrafted vs Grafted (right). See also Table S4.

**Table 1.** Summary mapping statistics by replicate group.

| Rootstock | Treatment | Mean Read Length (bp) | Total Read Length (bp) | Fold Coverage |
| --- | --- | --- | --- | --- |
| Ungrafted | Irrigated | 110 | 9,994,970,903 | 21 |
| Ungrafted | Unirrigated | 107 | 8,082,919,551 | 17 |
| Grafted | Irrigated | 110 | 9,328,021,049 | 20 |
| Grafted | Unirrigated | 108 | 10,548,329,716 | 22 |

The total level of methylation across clonal replicates of ‘Chambourcin’ varied only slightly by experimental group. UG vines displayed slightly greater genome-wide methylation than GR vines (Table 2). For both UG and GR vines, IR individuals displayed slightly greater genome-wide methylation than UI individuals. Genome-wide and across replicate groups, approximately 42.6%, 22.9%, and 4.6% of cytosines in the CpG, CHG, and CHH contexts, respectively, were methylated (H represents either A, C, or T) (Table 2; see also Table S1). Cytosine methylation detected in all sequence contexts in this study is similar to that typically observed in angiosperms, except the proportion of methylated cytosines in the CHH context was slightly greater than previously reported in grapevine (Niederhuth et al., 2016).

**Table 2.**
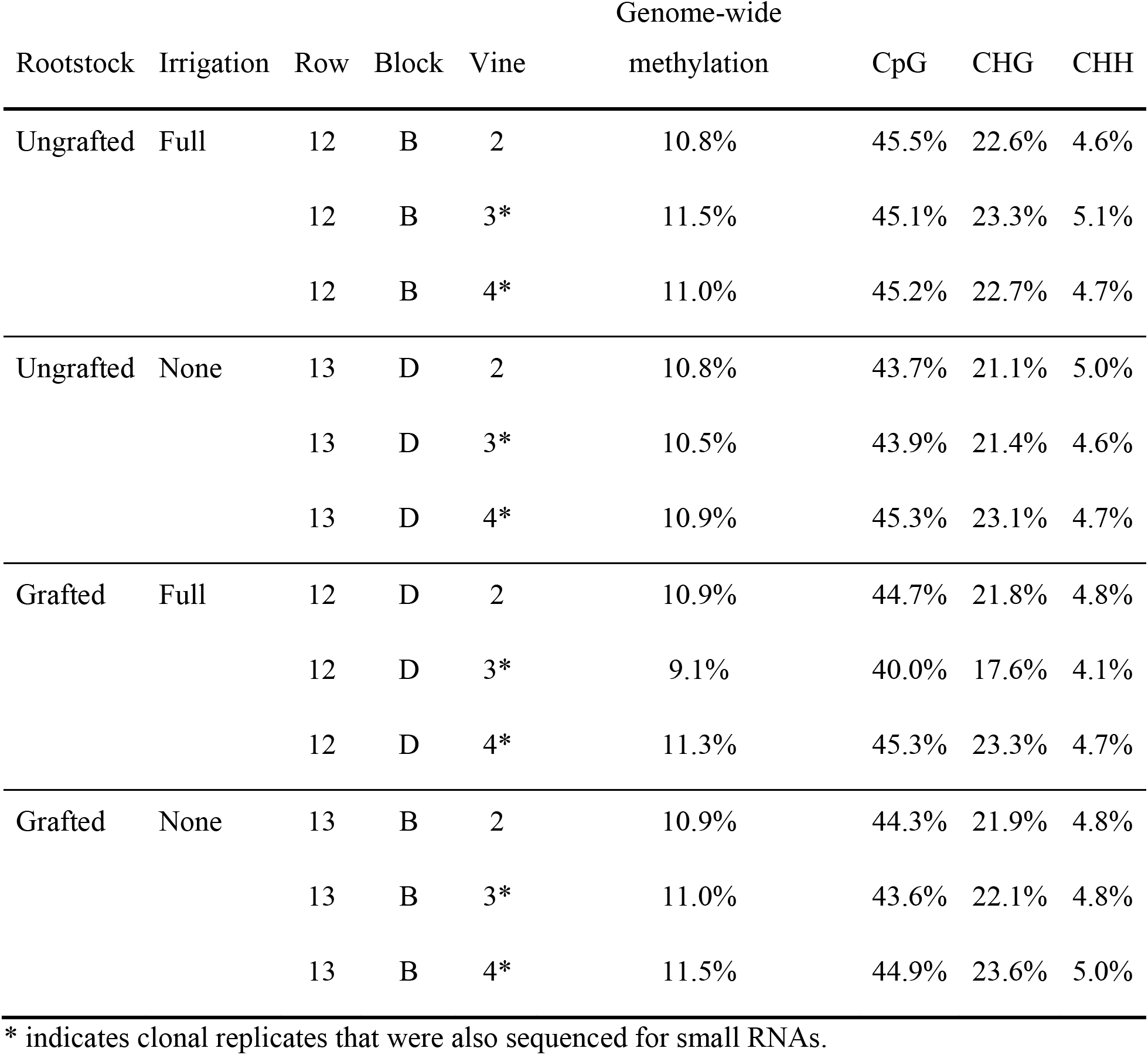
Estimated methylation levels by individual within replicate groups. * indicates clonal replicates that were also sequenced for small RNAs.

#### Genome-wide patterns of cytosine methylation reflect grafting and irrigation

PCA was used to visualize differences in methylation genome-wide among individuals. After filtering for regions averaging at least a 20% difference in methylation among individuals, we retained a dataset of 22,464 bp in the ~500Mb grapevine genome (see Table S2). PC1 and PC2 capture 21% and 17% of variation in methylation at sites among all vines (Fig. 1B). PC1 separated ungrafted (UG) individuals based on irrigation treatment, whereas grafted (GR) individuals in both irrigation treatments overlapped along PC1. PC2 separated grafted individuals from ungrafted individuals (Fig. 1B). This PCA provides strong support for methylation patterns differing by irrigation treatment in ungrafted vines, and by grafting condition.

#### Irrigated vs Unirrigated comparisons in ungrafted vines

To further investigate the extent to which irrigation treatment influenced variation in DNA methylation in ungrafted vines, we tested for significantly differentially methylated regions (DMRs; FDR adjusted *p*-value 0.05 in all subsequent analyses) in five categories: genome-wide, intragenic regions, and in CpG, CHG, and CHH sequence contexts. Because the classification of most annotated features have not been confirmed in grapevine, we will refer to these loci as “predicted features,” rather than “genes,” where appropriate. (For details on all DMRs, see Table S3.)

In UG vines, genome-wide DMRs were often more methylated in UI vines, whereas intragenic DMRs often displayed greater methylation in IR vines (Fig. 2). We identified 31 genome-wide DMRs between UG (IR vs UI) vines (Fig. 1C), a majority of which (24/31) were more methylated in UI conditions (Fig. 2). All genome-wide DMRs identified in UG vines were unique to this experimental group comparison. Next, we focused on intragenic methylation (methylation +/−2.5 kb predicted gene features), and identified two DMRs with increased methylation in IR conditions (Fig. 1D; Fig. 2).

**Figure 2.**
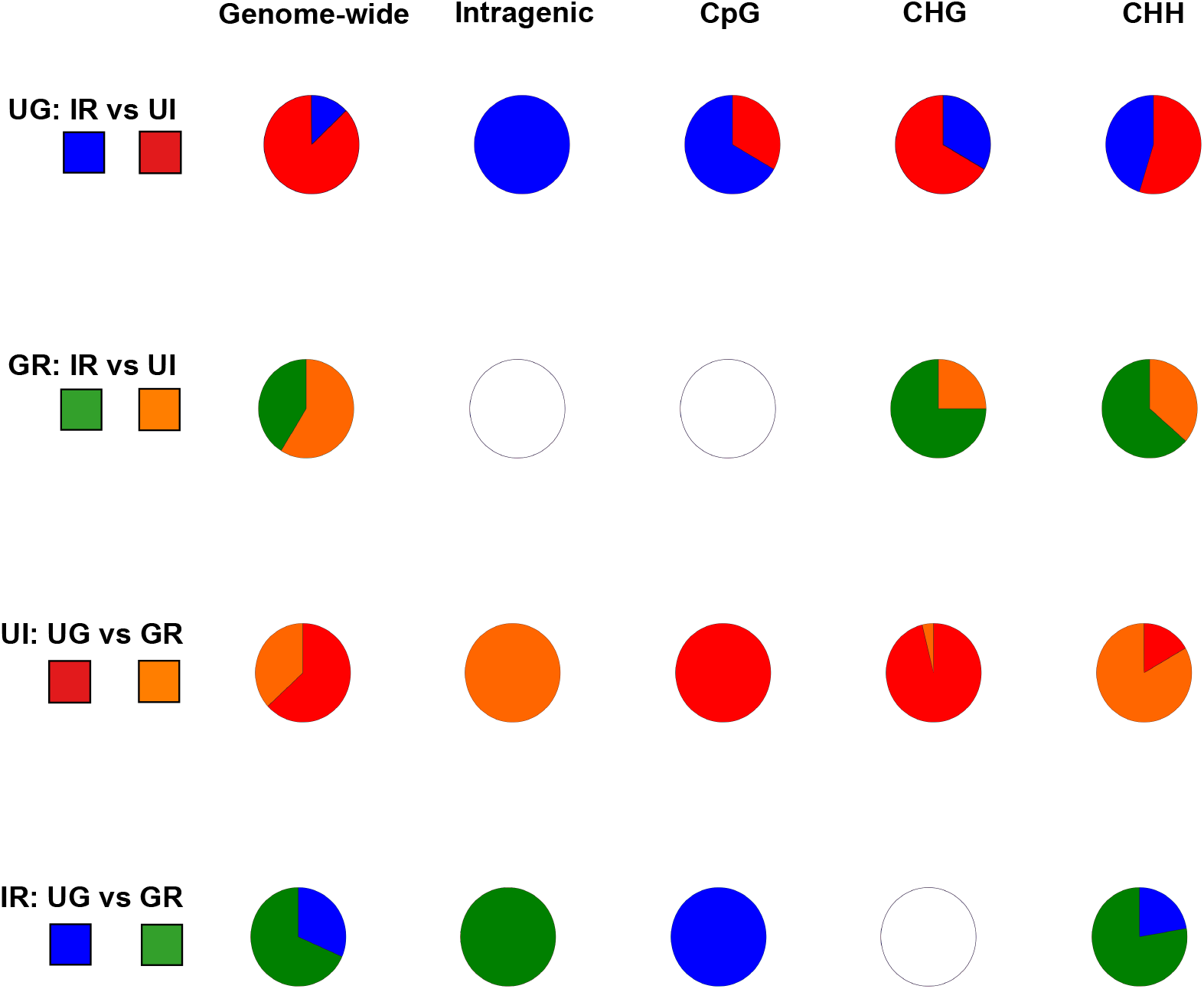
Proportion of DMRs in each category with elevated levels of methylation by experimental group comparisons. DMR categories are organized by column, experimental group comparisons are listed by row. Ungrafted, Irrigated vs Unirrigated: genome-wide, IR (7), UI (24); intragenic, IR (2), UI (0); CpG, IR (2), UI (1); CHG, IR (1), UI (2); CHH, IR (27), UI Grafted, Irrigated vs Unirrigated: genome-wide, IR (7), UI (10); intragenic, no DMRs; CpG, no DMRs; CHG, IR (3), UI (1); CHH, IR (21), UI (12). Unirrigated, Ungrafted vs Grafted: genome-wide, UG (12), GR (7); intragenic, UG (0), GR (2); CpG, UG (3), GR (0); CHG, UG (197), GR (7); CHH, UG (1), GR (4). Irrigated, Ungrafted vs Grafted: genome-wide, UG (6), GR (13); intragenic, UG (0), GR (1); CpG, UG (1), GR (0); CHG, no DMRs; CHH, UG (10), GR (35). Empty circles indicate no DMRs. See also Table S3.

In the analysis of patterns of CpG methylation (+/−2.5 kb over predicted gene features) in UG vines, we detected three DMRs (Fig. 1E), two of which demonstrated greater methylation in IR conditions (Fig. 2). DMRs in both CHG and CHH contexts frequently appeared within +/−2kb of predicted features and, contrary to methylation levels detected in CpG context, both CHG and CHH contexts often showed increased methylation in UI conditions (Fig. 2). Specifically, for CHG methylation, three DMRs were detected genome-wide and all were relevant to predicted features (Fig. 1F). Two of these CHG DMRs overlapped predicted features directly and had elevated methylation in UI conditions (Fig. 2). The other CHG DMR was located 378 bp upstream of a predicted feature and it was more methylated in IR conditions. The most DMRs in this comparison were detected in the analysis of CHH methylation; 61 DMRs were detected (Fig. 1G), and approximately half (30/62) were either directly overlapping a predicted feature or were located <2kb up- or downstream of a predicted feature. Of those 30 CHH DMRs, the majority (20/30 or 67%) directly overlapped predicted features; three were downstream and seven were upstream of predicted features. Out of the 61 total CHH DMRs, most (34/61) were more methylated in the UI vines (Fig. 2). Of CHH DMRs within +/− 2 kb of a predicted feature (30 DMRs), a majority (18/30) were also more methylated in UI vines.

In summary, DNA methylation patterns in ungrafted vines displayed a signature of irrigation in all DMR categories analyzed (genome-wide, intragenic, CpG, CHG, and CHH) (Fig. 1C-H). This irrigation effect was characterized by greater methylation in unirrigated conditions in three of the five DMR categories (genome-wide, CHG, and CHH) (Fig. 2). CHG and CHH methylation are not typically associated with genes. This suggests that most increases in methylation within UG vines occurred at non-genic regions when vines were unirrigated, which could be relevant to transposable elements. Notably, increased methylation surrounding predicted gene features in UG vines was more likely to occur under irrigated conditions. Therefore, the predicted genes associated with intragenic (+/−2.5 kb) and CpG DMRs would make ideal candidates for further investigation into how ungrafted ‘Chambourcin’ vines might be relying on the epigenome to moderate a transcriptional response to irrigation.

#### Irrigated vs Unirrigated comparisons in grafted vines

The effects of irrigation on patterns of DNA methylation were also compared in grafted vines. In contrast to what was observed in UG vines, a majority of DMRs in GR vines displayed elevated methylation in IR conditions (Fig. 2). Analysis of genome-wide methylation in GR (IR vs UI) vines identified 17 DMRs (Fig. 1C). Most (10/17) of which were more methylated in UI vines (Fig. 2). None of the genome-wide DMRs identified in GR vines were detected in UG vines. Zero intragenic DMRs were identified (Fig. 1D).

In GR vines, DMRs by sequence context displayed greater methylation in IR vines and non-genic regions experienced more changes in methylation than predicted genic regions. Analysis of patterns of CpG methylation (+/−2.5kb over predicted genes) detected zero DMRs (Fig. 1E). Analysis of CHG methylation revealed four DMRs (Fig. 1F), three of which were relevant to predicted features and were more methylated in IR conditions (Fig. 2). Two CHG DMRs overlapped predicted features directly and the other DMR was located 195 bp downstream of a predicted feature. In analysis of CHH methylation, 33 DMRs were detected (Fig. 1G) and the majority (21/33) were more methylated in IR conditions (Fig. 2). Approximately half (16/33) were either directly overlapping or <2kb of a predicted feature.

Relative to UG vines, GR vines displayed far less differential methylation between irrigation conditions; only three of five DMR categories (i.e., genome-wide, CHG, and CHH) displayed significant differences (Fig. 1C, F, G). Of those three significant DMR categories, we observed greater methylation in IR vines in the CHG and CHH sequences contexts, whereas UI conditions were characterized by a majority of DMRs with increased methylation genome-wide (Fig. 2). No DMRs were detected in GR vines under contrasting irrigation conditions in either the intragenic or CpG sequence context categories (Fig. 1D, E), suggesting that genic regions in ‘Chambourcin’ are perhaps not being targeted in methylome remodeling when vines are grafted, in spite of different environmental conditions.

#### Ungrafted vs Grafted in unirrigated conditions

To determine the effect of grafting on DNA methylation, we analyzed DMRs between UG and GR vines in a common irrigation treatment. First, we focused on genome-wide epigenomic variation under UI conditions; UG vines displayed more methylation than grafted vines for both genome-wide and intragenic DMRs in all sequence contexts (Fig. 1C, D; Fig. 2). We identified 19 genome-wide DMRs in all sequence contexts (Fig. 1C); 63% (12/19) were more methylated in UG vines than GR vines in unirrigated conditions (Fig. 2). Analyses exploring intragenic DMRs (+/−2.5 kb) in all sequence contexts identified two DMRs between UG and GR vines in UI conditions (Fig. 1D), both of which had elevated methylation in GR-UI vines (Fig. 2).

Between UG and GR vines in unirrigated conditions, sequence context specific DMRs were less consistent in methylation levels compared to the trend observed in genome-wide and intragenic DMRs (Fig. 2). When we searched CpG genic regions for differential methylation using a +/− 2.5 kb window, only three DMRs were detected (Fig. 1E). All three overlapped a gene and were more methylated in UG-UI vines (Fig. 2). Analysis of differential methylation in the CHG and CHH sequence contexts under unirrigated conditions showed the majority of CHG DMRs were more methylated in UG vines whereas most CHH DMRs were more methylated in GR vines. Most CHG and CHH DMRs overlapped a predicted gene feature. The analysis of CHG context-specific methylation identified 204 DMRs (Fig. 1F) of which 97% (197/204) were more methylated in UG vines (Fig. 2). Of the total CHG DMRs, 73% (148/204) either overlapped or were within 2kb of a predicted feature, of which 95% (141/148) were more methylated in UG vines (Fig. 2). Most genic CHG DMRs (130/141 or 88%) directly overlapped a predicted gene feature, accounting for 64% (130/204) of all CHG DMRs, whereas substantially fewer CHG DMRs were found upstream (15/148 DMRs) or downstream (7/148 DMRs) of predicted features. The search for differential methylation in the CHH sequence context revealed only five DMRs (Fig. 1G), three of which were overlapping or downstream of predicted features. Four showed greater methylation in GR-UI vines, including two that overlapped predicted features (Fig. 2). The final CHH DMR was located 855 bp downstream of a predicted feature and was more methylated in UG-UI vines.

In summary, patterns of differential methylation in unirrigated conditions demonstrated a general trend of higher methylation in UG vines across three of five total categories of DMRs (genome-wide, CpG, and CHG) (Fig. 2). The DMRs detected in the CHG sequence context in unirrigated conditions were particularly predominant (204) (Fig. 1F) and were overwhelmingly more methylated in UG vines (97%, with 64% directly overlapping predicted features) (Fig. 2). This is a potential focus of future investigation as previous research suggested that increased CHG methylation across genes can lead to the accumulation of CpG methylation over gene bodies, known as gene body methylation (gbM), which has been linked to the upregulation of some genes (Wendte et al., 2019).

#### Ungrafted vs Grafted in irrigated conditions

The final comparison focused on epigenomic variation between UG and GR vines that were irrigated (IR). In contrast to the unirrigated treatment, genome-wide analyses generally revealed greater methylation in GR vines than UG vines under irrigation (Fig. 2). We identified 19 genome-wide DMRs in all sequence contexts; 68% (13/19) were more methylated in GR vines (Fig. 1C). Intragenically, one DMR was identified in the analysis of all sequence contexts (Fig. 1D); it was also more methylated in GR vines (Fig. 2).

Patterns of methylation by sequence context were largely defined by differences in CHH methylation. Analysis of the CpG sequence context (+/−2.5kb of predicted gene), revealed only one DMR (Fig. 1E) and it was more methylated in UG vines (Fig. 2). In contrast to the prominent differences seen in CHG methylation in UI vines, zero CHG DMRs were detected between UG and GR vines that were IR (Fig. 1F). However, 46 CHH DMRs were found between UG and GR vines that received irrigation (Fig. 1G), 65% (30/46) of which either overlapped or were located within +/−2kb of a predicted feature. Overall, 80% (36/45) of CHH DMRs were more methylated in GR vines (Fig. 2) and 86% (26/30) of those located in or near a feature were more methylated in GR vines. Surprisingly, most of the feature-relevant CHH DMRs (14/30 or approx. 47%) were located within +/−2kb of a predicted feature, whereas 37% (11/30) and 16% (5/30) were found upstream and downstream of a feature, respectively. CHH DMRs that were more methylated in UG vines were located upstream of predicted features in all but one that overlapped a predicted feature. All five CHH DMRs located downstream of predicted features were more methylated in GR vines.

Overall, significant DMRs were detected between UG and GR vines that were irrigated in four out of five DMR categories. GR vines in irrigated conditions frequently showed greater methylation than UG vines (in three of the four DMR categories: genome-wide, intragenic (+/−2.5 kb), and CHH; Fig. 2). Generally, UG and GR vines that were irrigated displayed similar differences in genome-wide methylation relative to those detected under unirrigated conditions (Fig. 1A-G) except that the fewest DMRs detected in irrigated vines were in the CHG context and the most DMRs were detected in the CHH context (Fig. 1G). In the other three DMR categories, this comparison (UI: UG vs GR) revealed either few or a moderate amount of DMRs. Taken with our results that found significant differential methylation between UG and GR vines that were unirrigated, this suggests that grafting is a significant driver of DNA methylation differences in ‘Chambourcin’ vines.

### Characterization of small RNAs (sRNA)

#### Overview of sRNA

Small RNAs were characterized for eight of 12 clonal replicates used in BS-seq (two replicates in each of the four experimental treatments; see Supplemental Materials). As above, reads were mapped to the *Vitis vinifera* reference genome (PN40024 12Xv3). We conducted *de novo* cluster identification and analyses (quantitation and annotation), and generated read counts of sRNA for each sample (Table S5). The 21-nt size class of sRNAs (known as micro RNAs, or miRNAs, that can directly regulate gene expression) was most abundant, followed by the 24-nt class (known for their role in directing DNA methylation activities and targeting transposons for silencing) (Lewsey et al., 2016; Hardcastle et al., 2018). Less than 25% of all sRNAs mapped within predicted gene features (mRNA, gene bodies, and CDS). All sRNA loci intersected with methylated DNA in at least one sequence context in all experimental groups (Table S6).

#### Differential expression analysis of sRNA

The broader pattern detected in differential expression (DE) analysis of sRNA was similar to that observed in analyses of methylation. DE sRNAs varied both by irrigation and grafting treatment. Overall, very few significantly DE sRNA mapped to genes (only eleven sRNA, in total) (Fig. 1H; Table S4). In a comparison of UG and GR vines that were unirrigated, three significantly DE sRNA were detected. All three overlapped predicted genes and displayed higher expression in UG vines. A comparison of IR: UG vs GR vines again detected three significantly DE sRNA, which also overlapped predicted genes; these sRNA had higher expression in GR vines. There were no common DE sRNA between UI and IR conditions. DE sRNA were also found in a comparison of UG-IR vs UG-UI vines (Fig. 1H); five significantly DE sRNA were detected and overlapped predicted genes, two of which were more highly expressed in UG-IR vines and three showed higher expression in UG-UI vines. In contrast to our observations of DE sRNA in UG vines, we detected zero significantly DE sRNA in a comparison of GR-IR and GR-UI vines (Fig. 1H). The few DE sRNA detected in this study are consistent with published results from other work in grapevine (Zombardo et al., 2020).

## Discussion

In this study, we assessed methylome remodeling in 12 clonally replicated scions of the hybrid winegrape cultivar ‘Chambourcin.’ We observed that the ‘Chambourcin’ methylome is sensitive to both irrigation and grafting. Ungrafted vines showed a significant effect of irrigation treatment whereas grafted vines showed patterns distinct from ungrafted vines but with little effect of irrigation treatment. These results suggest that grafting might influence epigenomic sensitivity to environmental variation (e.g., water availability). We also detected differences in expression of sRNAs in response to the treatments that were consistent with the patterns seen in DNA methylation. These data indicate that irrigation and grafting affect the epigenome of a clonal grapevine.

### The Chambourcin methylome is plastic in response to irrigation treatment

The largest difference in methylation between ‘Chambourcin’ clones was measured in UG vines exposed to different irrigation treatments (Figs. 2 and 3). Some previous studies have detected effects of water availability on the epigenome of plants (Hubbard et al., 2014; Rico et al., 2014; Lafon-Placette et al., 2018; Li et al., 2020); however, a drought treatment did not alter methylation in *Arabidopsis* (Ganguly et al., 2017). Although we did not impose a drought treatment, the vines in this study experienced contrasting levels of irrigation for three growing seasons prior to when they were sampled, where half of the replicates received full replacement of evapotranspiration versus the other half which experienced periodic drought stress under natural conditions. Therefore, rather than reflecting an immediate cause-effect relationship between environment and methylation, differences in methylation that correspond to irrigation treatment in UG vines likely represent the cumulative effect of nearly three years of irrigation differences on the epigenome. The hypothesis for a gradual divergence in the methylome based on consistent pressure(s) from ecologically relevant factors is further supported by previous work that has identified an epigenomic signature of geographic provenance (Busconi et al., 2015; Dubin et al., 2015; Kawakatsu et al., 2016; Xie et al., 2017), as well as evidence that suggests this effect could be a form of epigenomically based memory (Avramova, 2015). The ability for a long-lived plant to maintain an epigenomic record in response to the long-term environmental conditions that it experiences might allow it to perform optimally in the specific environmental conditions in which it grows.

### The Chambourcin methylome bears a signature of grafting

The ‘Chambourcin’ methylome also bears a distinct signature of grafting, a result previously reported in studies in herbaceous plants (Wu et al., 2013; Cao et al., 2016; Kasai et al., 2016; Lewsey et al., 2016). The effect of the initial graft is traumatic to the plant (Melnyk, 2017) and elicits a multitude of genome-regulatory responses, but it also requires long-term management of molecular communication between different genomes, similar to what occurs in interspecific-hybrids. This result is concordant with previous research in different systems that has shown that genome-genome interactions result in an epigenomic signature, for example, in interspecific hybrids (Gaut et al., 2007; Landry et al., 2007; Ishikawa and Kinoshita, 2009; Wu et al., 2013; Cara et al., 2019) and in allopolyploids (Springer et al., 2015). One implication of the graft-associated epigenomic variation reported here is that specific scion genotypes could be epigenomically predictable based on the rootstock to which they are grafted. It would be interesting to investigate whether different rootstocks produce specific epigenotypes, whether rootstock-mediated epigenomic changes in the shoot are specific to the scion genotype or vary based on the scion, whether epigenomic modifications in the scion underlie predictable scion phenotypes, whether phenotypic stability is linked to epigenotypes generated by specific scion-rootstock combinations, and whether rootstock effects on scion methylation vary based on location. Furthermore, previous work in *Arabidopsis* has shown that it is possible for graft-inducible epigenomic modifications to be preserved from a grafted parent plant to seed offspring, which is linked to heritable phenotypes in the offspring (Virdi et al., 2015). If this is also true for woody clones, selection of desirable phenotypes from different combinations of grafted individuals in such species with long generation times could expedite the process of cultivar development in perennial crops. This is especially interesting to consider for wine grapes, where plant breeders might be able to enhance a cultivar’s terroir through epigenomically assisted selection of environmental inputs and clonal propagation of specific vines.

### The Chambourcin methylome shows a strong Irrigation × Grafting Interaction

Perhaps the most striking result of this study is in the interaction effect of irrigation and grafting on DNA methylation. The epigenome of ungrafted ‘Chambourcin’ exhibits a plastic response to contrasting irrigation conditions, but this epigenomic plasticity is constrained by the effect of grafting, observed as more consistent patterns of methylation in grafted vines. Although the practice of grafting in grapevine began as a way to protect grapevine roots from pests and pathogens, through its effect on the epigenome, grafting might provide additional viticultural benefits by stabilizing the vine’s phenotype and dampening its epigenomic response to different environmental conditions. This reduced epigenomic plasticity in a grafted scion might result in greater predictability of vine performance for agriculturally important traits like berry quality and yield in spite of environmental perturbations like variable precipitation. However, that grafting might act as a stabilizing or constraining force on a clonal grapevine scion’s epigenome exposed to different environments generates additional questions surrounding whether and how this might also alter a vine’s terroir, which should be the subject of additional study.

### Dynamic vs static DNA methylation in plants

Across the ‘Chambourcin’ genome, a small but significant portion of the methylome showed differences under experimental treatments (a total of 454 DMRs across the ~500Mb grapevine genome; Table S3). Although some studies have reported lability of DNA methylation in response to the environment, other studies have reported stability of DNA methylation in plants in variable environments (Niederhuth and Schmitz, 2014; Hagmann et al., 2015). We acknowledge that this is perhaps partially due to the challenge of identifying a small number of sites that differ in environmentally sensitive DNA methylation relative to the vast majority of stable methylation across the genome in plants (i.e., faithfully maintained within an individual and heritable between individuals). Another factor that may complicate the detection of environmentally responsive methylation is the biological noise present in plant DNA methylation data (van der Graaf et al., 2015), particularly since DNA methylation is a quantitative characteristic (measured as a proportionate value for each cytosine locus) and its measurement can be affected by the depth of sequencing achieved (Ziller et al., 2015). Another consideration is that DNA methylation can accumulate over time and cumulative epigenomic changes in clonal individuals exposed to consistent treatment(s) over multiple years (e.g., irrigation, grafting) might yield detectable signals that are not observed in shorter-term experiments. In the present study, we were able to detect an effect of experimental treatment on epigenomic variation by employing a single replicated genotype (i.e., clones), to which consistent treatments were applied over multiple years, and a sequencing design that ensured sufficient sequencing of replicates to achieve adequate depth of coverage and genome coverage. We recommend that future studies take these aspects of experimental design into account when investigating the effects of environment on variation in DNA methylation.

### Conclusions and Future Directions

In this study, we provide evidence for epigenomic plasticity in the grapevine clone ‘Chambourcin’: clonal replicates produce consistent, distinguishable variation in methylation patterns under different environmental treatments (irrigation) and agricultural practices (grafting). This epigenomic plasticity is likely to play an important role in the persistence of long-lived, clonal plants like grapevine; however, it is still largely unexplored in non-model systems and even less explored in clonal woody perennials in both natural and agricultural settings. Therefore, future work should focus on long-lived, clonal species in both natural populations and in orchards or vineyards. Natural experiments provide an opportunity to directly interrogate the establishment and structuring of epigenomic variation in nature, whereas the inclusion of established orchards and vineyards will allow researchers to explicitly investigate the capacity for genetically identical woody clones to respond epigenomically to environmental variation.

Currently, little is known regarding how epigenomic plasticity varies by genotype, how it is shaped in long-lived individuals under natural conditions, what role it plays in stress memory in perennials, and whether it responds when individuals are moved to common locations. An ideal study would track patterns of epigenomic variation in multiple clonal lineages growing in both natural populations and transplanted in a common garden. Such a design would provide an opportunity to differentiate among the effects of clonal genotype, geographic provenance, and the current environment on shaping patterns of epigenomic variation.

Furthermore, future work should also focus on how grafting might engage a plant’s epigenome, thereby altering or stabilizing the phenotype. An important area of inquiry will be whether the genetic identity of the clonal rootstock has a unique effect on the epigenome and/or phenotype of a scion, or if the act of grafting confers a generalized signature on the epigenome and, perhaps, phenotypic stability of a scion. Therefore, one target should be to answer questions such as: Can a signature in the methylome be detected by rootstock identity among multiple genetic lineages of rootstock? Or is there a signature in the methylome that is only detected in the scion at the resolution of a generalized grafting effect? And, how does this influence the stability or plasticity of a grafted scion’s epigenome when exposed to different experimental environments? By focusing on these types of questions and experiments we can improve our understanding of how epigenomic variation is generated and structured in diverse plant systems and what is its relationship to phenotypic variation and/or stability, as well as how plants respond to and remember their environment, all of which have broad implications for ecology, evolution, and agriculture.

## Methods

### Sample collection

Experimental grapevines located at the University of Missouri’s Southwest Research Center (Mt. Vernon, MO) were sampled just prior to berry harvest in September 2017. Vines were planted in 2009 and are either the ungrafted wine cultivar ‘Chambourcin’ or ‘Chambourcin’ grafted to rootstock 3309C (*Vitis riparia* x *V. rupestris*). Vines have been maintained in either full irrigation or no irrigation conditions since 2015. Twelve vines were sampled that represent four experimental treatment combinations (ungrafted, full irrigation; ungrafted, no irrigation; grafted, full irrigation; grafted, no irrigation) with three biological replicates per treatment group. The youngest leaf tissue on two shoots per vine was collected into cryovials and stored in liquid nitrogen for transport to a −80°C freezer.

### DNA extraction, bisulfite library construction, and sequencing

A modified CTAB protocol adapted from Azmat et al. (2012) was used to extract DNA from leaf tissue. DNA was quantified using a Qubit DNA High Sensitivity kit (Cat no. Q33231). Whole Genome Bisulfite Sequencing (WGBS) libraries were prepared using a ZR Pico Methyl-Seq™ Library Prep Kit (Cat no. 5456), then validated and quantified using a HS DNA kit on an Agilent 2100 BioAnalyzer. Libraries were pooled into three groups of four samples each and sequenced (2×150bp) on three lanes of Illumina HiSeq 4000 at Duke Center for Genomic and Computational Biology.

### WGBS quality filtering, read alignment, and methylation extraction

Bisulfite (BS) conversion rate was estimated by mapping sample reads against the *Vitis* chloroplast genome (see Supplemental Table S1). Quality filtering, alignment, deduplication, and methylation calling was performed in Bismark (Babraham Bioinformatics, Babraham Institute, Cambridge, UK; Krueger and Andrews, 2011). Vitis genome version GCF_000003745.3_12X was downloaded from NCBI. FastQC was performed before and after trimming. Reads were aligned and mapped in single-end mode and --non-directional option per Zymo recommendations. Reads were sorted in SAMtools before deduplication and methylation calling. Methylation information from both strands was combined using the --comprehensive option then output into context-specific files (CpG, CHG, and CHH). “coverage” files were generated with the --bedGraph option.

### Genome-wide methylation and identification of differentially methylated regions

DMRs were identified using SeqMonk (Babraham Bioinformatics, Babraham Institute, Cambridge, UK). SeqMonk uses the Ensembl genome annotation from the 12X, INSDC Assembly GCA_000003745.2 assembly (last modified 9/19/2016). Coverage files were imported into SeqMonk and replicates were grouped by experimental treatment. A running window approach (25,000 bp windows and steps), and read count quantitation identified coverage outliers 10 above the median to be filtered. Genome-wide cytosine methylation probes were created via running window (100 bp bins, 25 bp steps). In all DMR analyses, methylation over probes was assessed using the “Bisulfite methylation over features” pipeline, the mean value per probe reported, and Logistic Regression and EdgeR analyses were conducted and the results collated (multiple testing correction was applied; FDR cutoff *p*-value 0.05). A 15x minimum depth of coverage per replicate group was required in all analyses. Absolute methylation differences calculated pairwise between groups were: genome-wide ≥20%; intragenic ≥20%; CpG ≥10%; CHG ≥10%; CHH ≥10%. In all analyses, DMRs within 100 bp were grouped. Intragenic and CpG DMRs were defined as +/−2,500bp over predicted features. Analysis of CHG and CHH DMRs used the same running window probe generation protocols described above with the relevant methylation-by-sequence-context coverage files. For all DMR analyses, four comparisons were made: UG - IR vs UI; GR - IR vs UI; UI - UG vs GR; IR - UG vs GR.

### RNA extraction, library construction, and sequencing

RNA was extracted from the same leaf tissue as DNA using a ZR-Duet™ DNA/RNA MiniPrep Plus extraction kit (Cat. no. D7003). RNA samples were quantified using a Qubit RNA High Sensitivity kit (Cat no. Q32852). cDNA libraries were created from small RNA (sRNA) using an NEBNext® Multiplex sRNA Library Prep (set 1) kit (Cat no. E7300S/L). Libraries were validated and quantified on an Agilent 2100 BioAnalyzer System. Libraries were pooled and sequenced (1×50) on one Illumina Hiseq 4000 lane at Duke Center for Genomic and Computational Biology.

### Analysis of small RNA data

FastQC was performed on sRNA reads before and after trimming and filtering. Trim Galore! was used to carry out trimming and filtering of sRNA reads (minimum score of >Q20, maximum read length of 24-nt) (Martin, 2011). Alignment, annotation, and quantification of sRNAs was performed using ShortStack v3 (Axtell, 2013). Reads were mapped to *Vitis* genome version GCF_000003745.3_12X. The options specified were: zero mismatches, ‘unique’ multi-mapping alignment, 100 bp pads, minimum coverage designated in rpm, and strand identity determination. Count files from ShortStack were uploaded into SeqMonk; probes were generated via running window, “Percentile Normalisation Quantitation” using the 50^th^ percentile was used to calculate the additive scaling factor. Outlier Analysis filtered sRNAs at 10 above the median. Samples were grouped into “Replicate Datastores” by experimental group. Probes were created using “Read Position Probe Generator” (100 minimum reads, 1 valid position per window). sRNAs were filtered and reported to identify overlap with predicted genomic features. Probes were saved as annotation tracks and used as “features of interest” to identify areas of overlap with DNA methylation in each sequence context. DESeq2 was used to test for differential expression of sRNAs from un-normalized count files.

## Accession numbers

Sequence data from this article will be submitted to NCBI SRA under accession numbers XXX00000000 (WGBS) and XXX00000000 (sRNA-seq).

## Author Contributions

BW conducted the experiments and analyses, and wrote the paper. AM, MK, CE, and BW designed the experiments.

## Acknowledgements

The authors gratefully acknowledge the staff at the University of Missouri Southwest Research Farm and Keith Striegler who designed and established the Chambourcin experimental vineyard. We acknowledge members of the Miller Lab group and Edwards Lab group for careful comments on previous versions of the manuscript. Funding for this work was provided by Missouri Grape and Wine Institute Research Grant to AM, BW, CE, and MK, and NSF Plant Genome Research Program grant 1546869 to AJM and MK. No conflict of interest declared.

## Supplemental Information

Table S1. Summary of WGBS results and methylation estimates. Related to Table 2.

Table S2. Probe group report for genome-wide >/= 20% differential methylation. Related to Figure 1B.

Table S3. All DMRs identified by Logistic Regression and EdgeR analyses (FDR threshold 0.05). Related to Figure 1, C-G.

Table S4. Differentially expressed small RNA identified by DESeq2 from non-normalized count files. Related to Figure 1H.

Table S5. Summary of all putative small RNA identified in ShortStack v3.

Table S6. Intersection of small RNA loci and DNA methylation by sequence context. Related to Results “Overview of sRNA.”

